# Deletion of porcine *BOLL* causes defective acrosomes and subfertility in Yorkshire boars

**DOI:** 10.1101/2020.05.05.074724

**Authors:** Adéla Nosková, Christine Wurmser, Danang Crysnanto, Anu Sironen, Pekka Uimari, Ruedi Fries, Magnus Andersson, Hubert Pausch

## Abstract

A recessively inherited sperm defect of Finnish Yorkshire boars was detected more than a decade ago. Affected boars produce ejaculates that contain many spermatozoa with defective acrosomes resulting in low fertility and small litters. The acrosome defect was mapped to porcine chromosome 15 but the causal mutation has not been identified. We re-analyzed microarray-derived genotypes of affected boars and performed a haplotype-based association study. Our results confirmed that the acrosome defect maps to a 12.24 Mb segment of porcine chromosome 15 (P=3.38 × 10^−14^). In order to detect the mutation causing defective acrosomes, we sequenced the genomes of two affected and three unaffected boars to an average coverage of 11-fold. Read-depth analysis revealed a 55 kb deletion that segregates with the acrosome defect. The deletion encompasses the *BOLL* gene encoding the boule homolog, RNA binding protein which is an evolutionarily highly conserved member of the *DAZ* (deleted in azoospermia) gene family. Lack of *BOLL* expression causes spermatogenic arrest and sperm maturation failure in many species. Our study reveals that absence of *BOLL* is associated with a sperm defect also in pigs. The acrosomes of boars that carry the deletion in the homozygous state are defective suggesting that lack of porcine BOLL compromises acrosome formation. Our findings warrant further research to investigate the precise function of *BOLL* during spermatogenesis and sperm maturation in pigs.

## Main text

Reproduction in pigs is monitored at the population scale. Artificial insemination (AI) is widely practiced to enable high reproductive rates for genetically superior boars (Gerrits et al., 2005). Because each AI boar is mated to many sows, factors contributing to establish a pregnancy can be partitioned between males and females (Schaeffer, 1993). The heritability of boar fertility (i.e., the service-sire component) is relatively low, but the heritability of semen quality is higher (Wolf, 2009). Boar semen quality largely affects insemination success and litter size (Broekhuijse et al., 2011), (Holt et al., 1997).

The semen quality of AI boars is assessed immediately after ejaculate collection. Parameters that are monitored include concentration, morphology, vitality and motility of sperm. The rigorous monitoring of semen quality facilitates detecting and discarding ejaculates of insufficient quality. Boars that repeatedly produce ejaculates of insufficient quality may carry deleterious alleles that cause, e.g., spermatogenesis or sperm maturation defects (Sironen et al., 2006),(Sironen et al., 2011). Case-control association testing using dense microarray-derived genotypes may rapidly reveal genomic regions underpinning monogenic conditions (Charlier et al., 2008),(Bourneuf et al., 2017).

In 2010, Sironen et al. (Sironen et al., 2010) investigated an autosomal recessively inherited knobbed acrosome defect (KAD) in Finnish Yorkshire boars. Affected boars produced spermatozoa with had multiple acrosome aberrations including protruding granules and vacuoles. Depending on the proportion of sperm with defective acrosomes, affected boars were either infertile or their insemination success was low (Kopp et al., 2008). Apart from their reduced fertility, boars with the KAD were healthy. Case-control association testing between microarray (Illumina PorcineSNP60 Genotyping BeadChip) derived genotypes of affected and unaffected boars revealed that the KAD maps to an interval between 93 and 96 Mb on chromosome 15 (build 9 of the porcine genome). However, the underlying mutation had not been identified (Sironen et al., 2010).

To eventually identify the mutation causing the KAD, we re-analyzed the genotype data that were collected by Sironen et al. (Sironen et al., 2010). Following the re-assessment of the semen quality of affected boars, we retained 12 boars with the KAD as cases. Twenty-two boars from the Finnish Yorkshire breed that did not show the KAD were used as controls. All boars had been genotyped at 62,163 SNPs using Illumina PorcineSNP60 Genotyping Bead chips. The physical positions of the SNPs were according to the Sscrofa11.1 assembly of the porcine genome (Warr et al., 2019). Quality control was carried out using the plink software (version 1.9; (Chang et al., 2015)). We retained 42,212 autosomal SNPs that were genotyped in more than 90% of the samples, had MAF greater than 0.01 and did not deviate from Hardy-Weinberg proportions (*P*-value greater than 1 × 10^-4^). We imputed sporadically missing genotypes and inferred haplotypes using 25 iterations of the phasing algorithm implemented in beagle 5.0 (Browning et al., 2018), assuming an effective population size of 200 while all other parameters were set to default values. Subsequently, haplotype-based case-control association testing was implemented as described in (Venhoranta et al., 2014) and (Pausch et al., 2016). Specifically, we shifted a sliding window consisting of 20 contiguous SNPs corresponding to an average haplotype length of 1.08 ± 0.53 Mb along the autosomes in steps of 5 SNPs. Within each window, haplotypes with frequency greater than 1% were tested for association with the KAD using Fisher’s exact tests.

The association study revealed 35 haplotypes located between 66,900,647 and 102,308,093 bp on chromosome 15 (SSC15) that exceeded the Bonferroni-corrected significance threshold (5.35 x 10^-7^). The most significantly associated haplotype (P=3.38 x 10^-14^) was located between 100,183,707 and 101,401,120 bp (Figure 1A). We did not detect haplotypes on chromosomes other than SSC15 that were significantly associated. The 12 boars with the KAD shared a 12.24 Mb segment (between 90,012,606 and 102,260,926 bp) of extended homozygosity (Figure 1B) that was not detected in the 22 control boars. According to the Refseq (version 106) annotation of the porcine genome, this segment encompasses 51 genes including *STK17b* and *HECW2*, i.e., two genes that were prioritized as positional candidate genes by Sironen et al. (Sironen et al., 2010).

**Figure 1.**
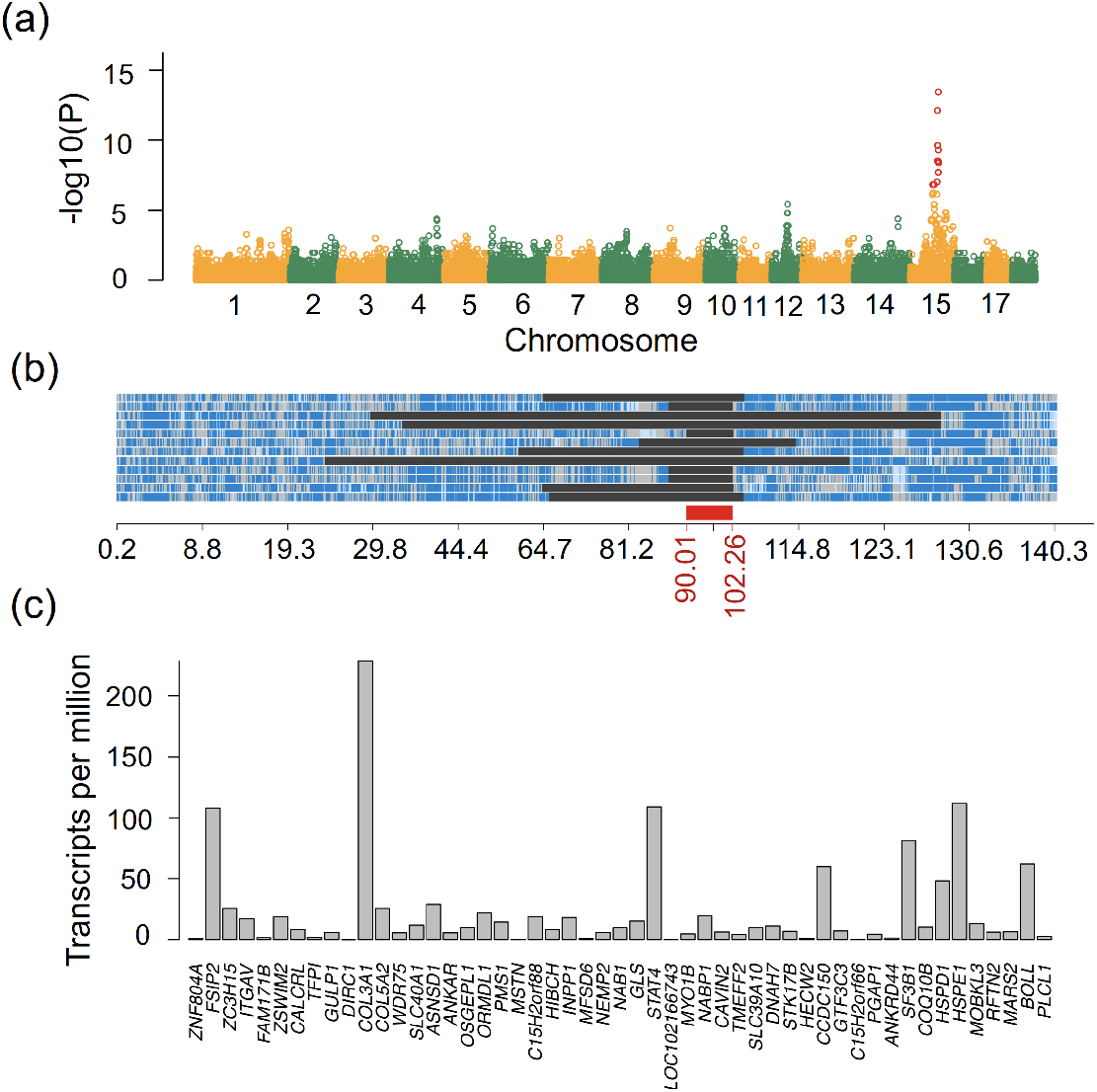
A knobbed acrosome defect in Finnish Yorkshire boars maps to a 12.24 Mb segment on porcine chromosome 15. (a) Manhattan plot of a haplotype-based genome-wide association study. Red color represents 35 haplotypes with a *P* value less than 5.35 x 10^-7^ (Bonferroni corrected-significance threshold). (b). Homozygosity mapping in 12 boars with defective acrosomes. Blue and pale blue represent homozygous genotypes (AA and BB), heterozygous genotypes are displayed in light grey. Dark grey areas represent segments of extended homozygosity. The red bar indicates a 12.24 Mb segment of extended homozygosity that was shared in all affected boars. (c) mRNA abundance of 51 genes that are located within the segment of extended homozygosity in testis tissue of pubertal boars.

We hypothesized that the mutation causing defective acrosomes might affect a transcript that is abundant in porcine testis. In order to quantify the mRNA abundance of genes within the segment of extended homozygosity, we investigated total RNA sequencing data of testis tissue from seven pubertal Piétrain and Landrace boars that were collected by Robic et al. (Robic et al., 2019). We used the kallisto software (Bray et al., 2016) to pseudo-align the RNA sequencing reads to the porcine transcriptome (Refseq version 106) and quantify transcript abundance. Seven genes within the 12.24 Mb segment *(FSIP2, COL3A1, STAT4, CCDC150, SF3B1, HSPE1, BOLL)* were highly expressed (transcripts per million (TPM) > 50) in the testes (Figure 1C). Among these genes, the expression of *FSIP2* and *BOLL* is germ cell-specific and mutations within both genes cause defective sperm flagella, spermatogenic failure and male infertility (Brown et al., 2003),(Martinez et al., 2018),(Shah et al., 2010). The two positional candidate genes prioritized by (Sironen et al., 2010) were lowly expressed *(STK17b:* TPM=6.9, *HECW2:* TPM=1.2).

Next, we sequenced two affected boars (SAMEA6798259, SAMEA6798260), one heterozygous haplotype carrier (SAMEA6798263) and two boars (SAMEA6798261, SAMEA6798262) that did not carry the KAD-associated haplotype to 11.1-fold sequencing read depth using 2 ×126 bp paired-end reads. Illumina TruSeq DNA PCR-free libraries with 350 bp insert sizes were prepared and sequenced with an Illumina HiSeq2500 instrument. We used the fastp software (Chen et al., 2018) to remove adapter sequences and reads that had phred-scaled quality less than 15 for more than 15% of the bases. The filtered reads were aligned to the Sscrofa11.1 assembly of the porcine genome using the mem algorithm of the bwa software (Li, 2013). We marked duplicates using the picard tools software suite (https://github.com/broadinstitute/picard) and sorted the alignments by coordinates using sambamba (Tarasov et al., 2015). Sequence variants (SNPs and Indels) of the five sequenced animals were genotyped together with 93 pigs from breeds other than Finnish Yorkshire for which sequence data were available from our in-house database using the multi-sample variant calling approach of the genome analysis toolkit (gatk, version 4.1.0; (Depristo et al., 2011)). We filtered the sequence variants by hard-filtering according to best practice guidelines of the gatk. Functional consequences of polymorphic sites were predicted according to the Refseq annotation (version 106) of the porcine genome using the variant effect predictor tool from Ensembl (McLaren et al., 2016).

In order to detect plausible candidate causal mutations for the KAD, we considered 78,861 SNPs and Indels that were detected within the 12.24 Mb segment of extended homozygosity on chromosome 15. We retained variants that were homozygous for the alternate allele in two affected boars, heterozygous in the haplotype carrier and homozygous for the reference allele in two boars that did not carry the KAD-associated haplotype. This filtration revealed 89 variants that were compatible with recessive inheritance. However, none of them was a compelling candidate causal mutation because all were located in either intergenic or intronic regions. In order to investigate if large deletions or duplications might underpin the KAD, we inspected sequencing read depth within the segment of extended homozygosity on SSC15 using the mosdepth software (Pedersen and Quinlan, 2018). The two boars with the KAD had virtually no reads mapped to an ~55 kb interval suggesting that they were homozygous for a deletion (Figure 2). The boar that carried the top-haplotype in the heterozygous state carried the deletion in the heterozygous state (Figure 2d). The deletion was absent in two boars that did not carry the KAD-associated haplotype and in 93 control animals from breeds other than Finnish Yorkshire. Defining the precise boundaries of the deletion was difficult because the mapping quality of the sequencing reads flanking the deletion was low (Supporting File 1). Results of the basic local alignment search tool (blast) indicated that the flanking regions contain multiple species-specific repeats that likely complicated the accurate alignment of reads and also prevented the design of specific primers for PCR amplification. Based on read depth analyses and alignment visualization, we determined the boundaries of the deletion at approximately 101,549,770 and 101,604,750 bp (Supporting File 1).

**Figure 2:**
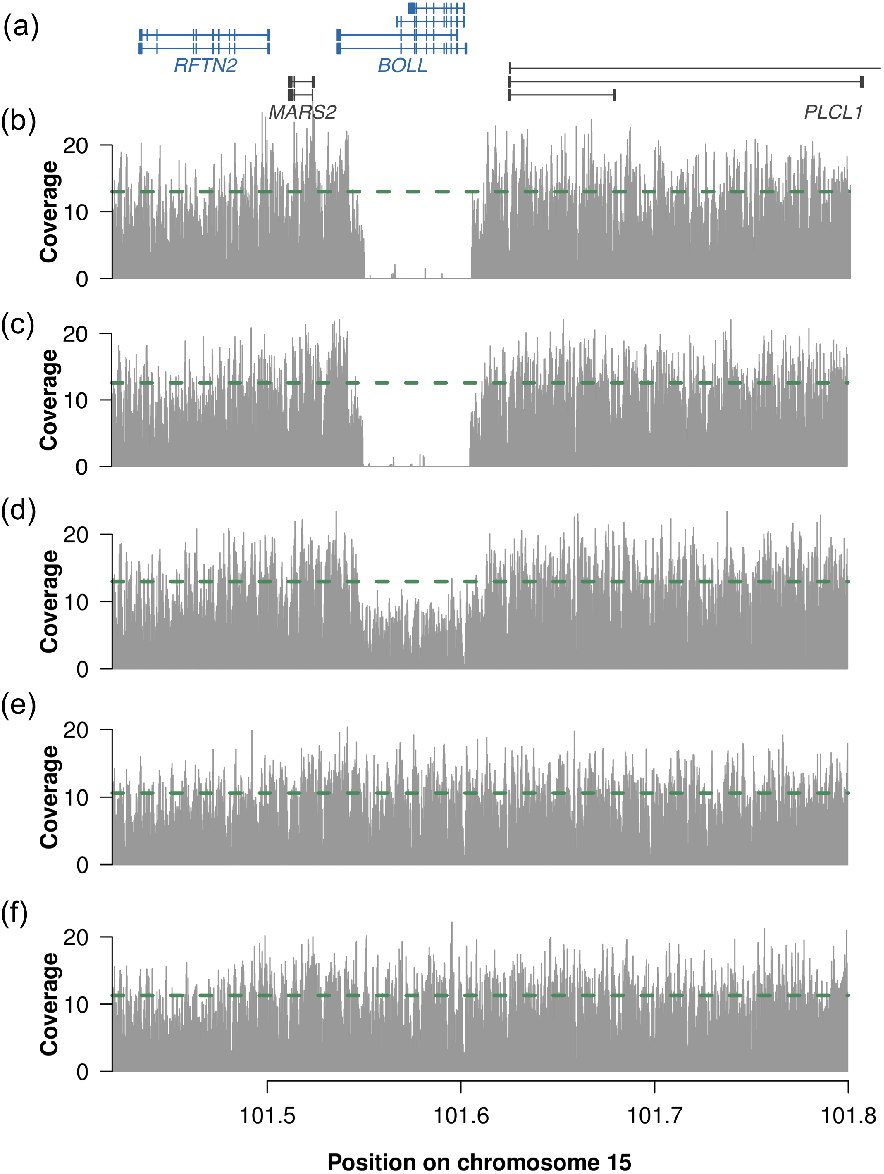
A 55 kb deletion segregates with the knobbed acrosome defect. Sequence read depth averaged over 250 bp windows for two homozygous (b, c), one heterozygous (d) and two control boars (e, f) at between 101.4 and 101.8 Mb on porcine chromosome 15. The green dashed line represents the average sequence read depth for the five boars. The two boars with the knobbed acrosome defect (a, b) have virtually no coverage between 101,549,770 and 101,604,750 bp. The coverage of the heterozygous boar in that region (d) drops to approximately half the average coverage. Blue and grey color represent genes and their isoforms located on the forward and reverse strand, respectively (a).

The 55 kb deletion encompasses only the *BOLL* gene encoding the boule homolog, RNA binding protein which is an evolutionarily conserved ancestral member of the *DAZ* (deleted in azoospermia) gene family (Figure 2a). The expression of *BOLL* mRNA is essential for male reproduction in many metazoan species (Shah et al., 2010). We also found that *BOLL* mRNA is highly abundant (TPM=62) in porcine testes suggesting that it may also be required for male reproduction in pigs (Figure 1C). Loss-of-function alleles in mammalian and insect orthologs of *BOLL* cause male-specific infertility resulting from absence of spermatozoa due to meiotic arrest (Eberhart et al., 1996),(Xu et al., 2003),(Shah et al., 2010),(Sekiné et al., 2015). It is very likely that the 55 kb deletion and resulting absence of *BOLL* expression causes the sperm defect of the Finnish Yorkshire boars. Our findings and the investigations of Sironen et al. (Sironen et al., 2010) and Kopp et al. (Kopp et al., 2008) suggest that lack of porcine *BOLL* is associated with impaired sperm maturation. The sperm count in ejaculates of boars with the KAD was not reduced (Kopp et al., 2008), suggesting that absence of *BOLL* does not notably affect meiotic progression in pigs. Thus, our findings corroborate that the role of *BOLL* during spermatogenesis and sperm maturation may vary between species (Li et al., 2019) and warrant further research to elucidate its precise function in pigs.

In conclusion, we report a candidate causal mutation for a sperm disorder of Finnish Yorkshire boars. A 55 kb deletion encompassing porcine *BOLL* is very likely causal for defective acrosomes and low male fertility. Once more, our findings highlight that revisiting unresolved genetic disorders with whole-genome sequencing technologies may readily reveal candidate causal mutations that remained undetected so far (Fang et al., 2020).

## Acknowledgements

We acknowledged financial support from SUISAG, Micarna SA, the ETH Zürich Foundation and the Finnish Veterinary Research Foundation.

## Availability of data

Whole-genome sequence data of five Yorkshire boars are available at the European Nucleotide Archive (ENA) of the EMBL at the BioProject PRJEB37956 under sample accession numbers SAMEA6798259 – SAMEA6798263. RNA sequencing data of testicular tissue samples of seven pubertal boars are available at the European Nucleotide Archive (ENA) of the EMBL at the BioProject PRJNA506525 under sample accession numbers SAMN10462191 - SAMN10462197. Whole-genome genotyping data of affected and unaffected boars are available through (Sironen et al., 2010).

---

**Supporting File 1:**
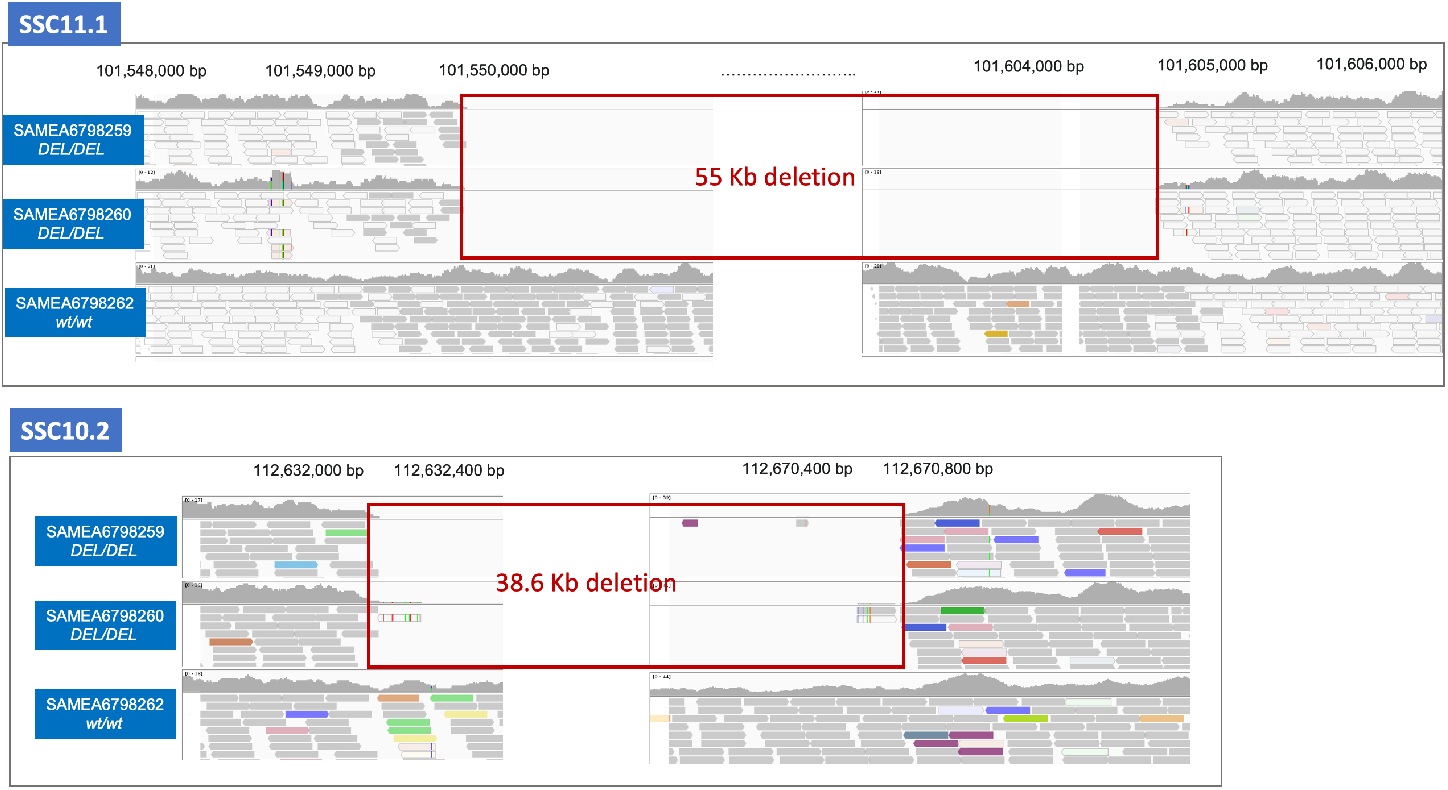
A large deletion segregates with the KAD-associated haplotype. Screen captures of igv outputs from whole genome sequence alignments of two boars homozygous for the deletion (SAMEA6798259, SAMEA6798260) and one boar homozygous for the corresponding reference nucleotides (SAMEA6798262). The length of the deletion is ~55 and ~38.6 kb according to the SSC11.1 (upper panel) and SSC10.2 (lower panel) assembly of the porcine genome. Please note that all alignments flanking the deletion are displayed with light grey borders and transparent fill (indicating low mapping quality).

